# Polyploidy and elevation contribute to opposing latitudinal gradients in diversification and species richness in lady ferns (Athyriaceae)

**DOI:** 10.1101/351080

**Authors:** Ran Wei, Richard H. Ree, Michael A. Sundue, Xian-Chun Zhang

**Affiliations:** State Key Laboratory of Systematic and Evolutionary Botany, Institute of Botany, the Chinese Academy of Sciences, Beijing 100093, China; Life Sciences Section, Integrative Research Center, The Field Museum, Chicago, IL 60605, USA; The Pringle Herbarium, University of Vermont, 27 Colchester Drive, Burlington, VT 05405, USA

**Keywords:** Athyriaceae, elevation, latitudinal gradient, macro-evolution, SSE, models, polyploidy

## Abstract

- In ferns, the temperate-tropical sister clades *Athyrium* and *Diplazium* present an opportunity to study a latitudinal contrast in diversification dynamics.
- We generated a taxonomically expanded molecular chronogram and used macroevolutionary models to analyze how diversification rates have changed through time, across lineages, and in concert with changes in elevation and ploidy. We tested a novel model of cladogenetic state-change in which polyploidy can arise as an infraspecific polymorphism, with diversification parameters distinct from those of pure diploids and polyploids.
- Both *Athyrium* and *Diplazium* accelerated their diversification near the Oligocene-Miocene transition. In *Diplazium*, the rate shift is older, with subsequent net diversification somewhat slower and suggestive of diversity-dependence. In *Athyrium*, diversification is faster and associated with higher elevations. In both clades, polyploids have the highest rate of net accumulation but lowest (negative) net diversification, while the converse is true for polymorphic species; diploids have low rates of both net accumulation and diversification.
- Diversification in *Athyrium* may have responded to ecological opportunities in expanding temperate habitats during the Neogene, especially in mountains, while the pattern in *Diplazium* suggests saturation in the tropics. Neopolyploids are generated rapidly, primarily through accelerated cladogenesis in polymorphic species, but are evolutionary dead ends.

## Introduction

How does the tempo (rate) and mode (process) of lineage diversification vary across latitude? Consideration of this question has largely focused on explaining the latitudinal gradient in species richness, especially with respect to how temperate-tropical contrasts in time-integrated area, temperature and metabolic rate, and ecological opportunity might influence the rate of speciation (reviewed in Schluter, 2015). A general prediction is that, all else being equal, tropical clades should be older, and/or have higher net diversification, compared to temperate clades (Wiens & Donoghue, 2004; Mittelbach *et al.*, 2007). In this context, temperate-tropical sister clades that otherwise share similar ecologies offer opportunities to study the correlates and potential drivers of diversification across latitudes over a common timescale.

One such opportunity presents itself in the lady-fern family Athyriaceae, in which the primarily temperate genus *Athyrium*, including allied genera, is sister to the primarily tropical *Diplazium* (Wei *et al.*, 2013, 2018). These clades respectively include about 220 and 350 species that occur in habitats that range from wet and shaded streambanks at lower altitudes to dry and open grasslands higher up (Tryon & Tryon, 1982; Wang *et al*., 2013). Both clades are more or less globally distributed, with the majority of species in tropical and temperate Asia (Wang *et al.*, 2013) and the tropical Andes (Tryon & Tryon, 1982), and each has been the subject of recent systematic and biogeographic studies (Wei *et al.*, 2013, 2015, 2018). Here, we generate a more comprehensive dataset for comparative analyses of diversification: in particular, how the timing, extent, and phylogenetic distribution of rate heterogeneity differ in these clades, and what this reveal about their responses to potential drivers of diversification.

In considering such drivers, a common theme that emerges from prior studies of fern diversification is ecological opportunity. Seminal work at broad taxonomic and deep temporal scales emphasized how fern lineages, particularly epiphytes, proliferated ‘in the shadow of angiosperms’, i.e. in novel habitats created as the latter ascended in the late Cretaceous/early Cenozoic (Schneider *et al.*, 2004; Schuettpelz & Pryer, 2009; but see Testo & Sundue, 2016). More recently, birth–death models fitted to phylogenies and the fossil record showed evidence for diversity-dependent origination, interpreted as reflecting opportunistic niche-filling, and extinction driven by environmental change (Lehtonen *et al.*, 2017). For example, in Polypodiaceae, faster diversification is correlated with changes in elevation, a proxy for habitat type, suggesting that niche shifts drive lineage proliferation (Sundue *et al.*, 2015). In the case of *Athyrium*-*Diplazium*, one might predict that diversification in *Athyrium* was accelerated by ecological opportunities arising during the expansion of temperate habitats during the Neogene, compared to *Diplazium*, which might be in the later stages of diversity-dependent clade growth, reflecting its persistence in more continuously stable tropical environments. Both clades might show responses to elevation, reflecting their respective colonization of temperate and tropical mountains, such as temperate Hengduan Mountains and the tropical Kinabalu and Andes.

Another potential driver of diversification that is not directly related to latitude, but demands attention due to its prevalence in ferns, is polyploidy (e.g. Schneider *et al.*, 2017). There is evidence that more than 50% of the species in the *Athyrium-Diplazium* clade are polyploid (e.g. Praptosuwiryo, 2008; Bir & Verma, 2010). The effect of polyploidy on plant diversification has received much attention from both theoretical and empirical perspectives (e.g. Mayrose *et al.*, 2011; Soltis *et al.*, 2016; reviewed in Vamosi *et al.*, 2018). In particular, polyploid formation as a mode of speciation motivated the development of cladogenetic state-dependent diversification models, in which the rate of speciation coincident with state change is parameterized (Mayrose *et al.*, 2011; Zhan *et al.*, 2016; see also Goldberg & Igić, 2012). These have since been used to show that neopolyploids tend to be evolutionary 'dead ends' (Mayrose *et al.*, 2011; Arrigo & Barker, 2012) due to effects such as gene imbalance of the sex chromosomes during the meiosis (Orr, 1990), increased abortion in heteroploid hybrids (Ramsey & Schemske, 2002), and inefficiency of selection for multi-copy genes in polyploids (Wright, 1969). We wish to know if this holds true in the diversification dynamics of *Athyrium* and *Diplazium*.

Previous studies of neopolyploidy using cladogenetic state-change models only considered two states for species, diploid and polyploid (Zhan *et al.*, 2016). However, infraspecific variation in ploidy is common in plants (Wood *et al.*, 2009); in the *Athyrium-Diplazium* clade, 5–10% of species have polyploid individuals recorded within one or more diploid populations (Walker, 1966; Tryon & Tryon, 1982; Takamiya *et al.*, 1999, 2000; Takamiya & Ohto, 2001; Praptosuwiryo, 2008; Bir & Verma, 2010). Such cases of polymorphism may represent incipient speciation via polyploidization. To test this idea, we constructed cladogenetic state-change models that include three states, diploid, polyploid, and a polymorphic diploid/polyploid state, with free parameters assigned to the rate at which polymorphic species give rise to new neopolyploids via cladogenesis events.

Our study can be summarized as a temperate-tropical contrast in diversification dynamics, focusing on how sister clades have responded to common potential drivers of diversification. Specifically, our aims are to reconstruct a time-scaled global phylogeny of the *Athyrium-Diplazium* clade, estimate variation in diversification rates through time, and infer the effects of elevation and polyploidy. Our intent is to shed light on latitudinal differences in the tempo and mode of diversification, and contribute to a more general understanding of the latitudinal diversity gradient.

## Materials and Methods

### Taxon sampling and molecular data

Our sampling strategy followed the most recent systematic studies of Athyriaceae (Wei *et al.*, 2013, 2015, 2018; PPG I, 2016). In this study, the circumscription of *Athyrium* includes four closely related genera (*Anisocampium*, *Athyrium*, *Cornopteris* and *Pseudathyrium*) and ten sections (sects. *Athyrium*, *Biserrulata*, *Dissitifolia*, *Mackinnoniana*, *Otophora*, *Polystichoides*, *Rupestria*, *Spinulosa*, *Wallichiana* and *Yokoscentia*) (Supporting Information Table S1). Since no sectional classification in the *Diplazium* clade (especially in subgenus *Callipteris*), we tentatively subdivided *Diplazium* into 11 clades (CHI, DIP, LEP, MET, MON, NEO, PSE, SEA, SIB, SPE and UNK) according to their morphological similarity, phylogenetic relationship and biogeographical affinity (Table S1).

In total, 85 out of *c.* 220 species of *Athyrium* (39%) and 129 out of *c*. 350 species of *Diplazium* (37%) on the global scale were sampled, including representatives from all taxonomic subdivisions or infrageneric groups covering the entire geographical distribution range and ecological habitats of these two genera. Voucher information is listed in Table S2. We included 26 representatives from Aspleniaceae, Athyriaceae, Blechnaceae, Cystopteridaceae, Desmophlebiaceae, Diplaziopsidaceae, Dryopteridaceae, Hemidictyaceae, Onocleaceae, Rhachidosoraceae, Thelypteridaceae and Woodsiaceae as outgroups and to provide appropriate nodes for fossil calibrations.

We assembled a dataset of eight chloroplast regions (*atp*A, *atp*B, *mat*K, *rbc*L, *rpl*32-*trn*P, *rps*4, *rps*4-*trn*S, *trn*L-F) with a total aligned length of 8297 bp. We used 1129 previously published sequences and generated 336 new sequences (Table S2). DNA extraction, amplification, sequencing, and alignment protocols followed Wei *et al*. (2013, 2018).

### Divergence time estimation

We inferred the posterior distribution of chronograms using BEAST 1.8.4 (Drummond & Rambaut, 2007) with the alignment partitioned by protein coding regions and intergenic spacers. Preliminary analyses rejected a strict molecular clock, so we used the uncorrelated lognormal relaxed clock model with a birth–death prior on tree shape. Eight independent runs of 30 million generations, sampling every 1000 generations, were carried out on the Cyberinfrastructure for Phylogenetic Research (CIPRES) Science Gateway (http://www.phylo.org; Miller *et al.*, 2010). The resulting log files were combined (with the first 50% samples discarded as burn-in) using LOGCOMBINER 1.8.4 (Drummond & Rambaut, 2007) and checked in TRACER 1.6 to make sure the effective sampling sizes for most of the relevant estimated parameters were well above 200. The tree files were resampled from each run using the same burn-in strategy in LOGCOMBINER. The maximum clade credibility (MCC) topology with a posterior probability limit of 0.5 and mean branch lengths was summarized using TREEANNOTATOR 1.8.4 (Drummond & Rambaut, 2007).

Four calibration points were used. For the root node the prior was a Normal distribution with a mean age of 107.29 Myr and a SD of 10 (with 95% highest posterior density [HPD]: 90.84–123.7 Myr) to cover the range of published split times from 91 Myr to 125 Myr including the 95% HPD intervals (Schneider *et al.*, 2004; Schuettpelz & Pryer, 2009; Rothfels *et al.*, 2015; but see Testo & Sundue, 2016). We used the Paleocene fossil of *Onoclea* (Rothwell & Stockey, 1991) to constrain the minimum age of the Onocleaceae crown group with a lognormal prior distribution with an offset of 54.5 Myr and mean of 1.0 (with 95% HPD: 55.02–68.58 Myr). We used earliest fossil record of *Woodwardia*, known from Paleocene deposits of North America, to constrain the crown node of Blechnaceae (Collinson, 2001), using the same prior settings. We assigned the *Diplazium*-like fossil reconstructions (*Makotopteris princetonensis*, Stockey *et al.*, 1999; *Dickwhitea allenbyensis*, Karafit *et al.*, 2006) found in the Middle Eocene deposits of British Columbia in North America to the stem lineage of *Diplazium* (Wei *et al.*, 2015), yielding a lognormal prior with an offset of 36.5 Myr and mean of 2.0 (with 95% HPD: 37.93–74.78 Myr) for the crown node of the *Athyrium-Diplazium* clade.

### State-independent diversification analysis

We used Bayesian Analysis of Macroevolutionary Mixtures (BAMM) 2.2.2 (Rabosky, 2014) to estimate rates of speciation (λ) and extinction (μ) through time and across clades on the MCC tree, pruned to the ingroup *Athyrium-Diplazium* clade. To account for incomplete sampling, we estimated the sampling fractions of infrageneric groups (Table S1) from our own taxonomic knowledge and the literature (e.g. Ching, 1964; Kato, 1977; Tryon & Tryon, 1982; Hsieh, 1986; Tryon & Stolze, 1991; Zhang, 1992; Stolze *et al.*, 1994; Wang, 1997; Mickel & Smith, 2004; Y.C. Liu *et al.*, 2011; Wang *et al.*, 2013; Wei *et al.*, 2013, 2018). Four MCMC analyses were run for 3 million generations each, sampled every 1000 generations. Each run was checked to ensure that the effective sample size (ESS) exceeded 200, and the first 10% of samples were discarded as burn-in. Results were summarized using BAMMTOOLS (Rabosky *et al.*, 2014). To evaluate the best model generated by BAMM (compared with a null model with no diversification rate shifts), we relied on Bayes Factors calculated with the COMPUTEBAYESFACTOR function of BAMMTOOLS.

### State-dependent diversification analyses

We tested two variables for an association with diversification: mean elevation and ploidy. Data were obtained from a variety of sources, including online databases (Global Plants, https://plants.jstor.org; Tropicos, http://www.tropicos.org; IPCN, http://www.tropicos.org/Project/IPCN; CCDB, http://ccdb.tau.ac.il/Pteridophytes/Athyriaceae), herbarium specimens, and cytological reports (e.g. Mehra & Bir, 1960; Walker, 1966, 1973; Tryon & Tryon, 1981; Takamiya *et al.*, 1999, 2000; Takamiya & Ohta, 2001; Praptosuwiryo, 2008; Bir & Verma, 2010), and are available in Table S3.

To test the hypothesis of net diversification accelerated by divergence in elevation (Sundue *et al.*, 2015), we used BAMM to estimate the rates of elevation change (β) at the tips of the MCC tree, and performed phylogenetic generalized least squares (PGLS) regression of β on tip values of *r* = λ – μ using GEIGER (Harmon *et al.*, 2008). In addition, we tested for a direct effect of elevation using the quantitative-state speciation-extinction (QuaSSE) model (FitzJohn, 2010). We considered seven candidate relationships between elevation (log-transformed) and λ: 1) λ is constant and independent of elevation; 2) linear; 3) sigmoid; 4) unimodal, represented by a vertically offset Gaussian function; and another three models (linear, sigmoid, and unimodal) with a directional tendency (Table S4). We ran each model on the entire ingroup clade as well as separately on *Athyrium* and *Diplazium* using DIVERSITREE 0.9-10 (FitzJohn, 2012). We compared models using the Akaike information criterion (AIC).

To explore the effect of ploidy on diversification we used the cladogenetic state change speciation-extinction (ClaSSE) model (Goldberg & Igíc, 2012). Species were scored as 1 (polymorphic, i.e. diploid with records of intermingled polyploids), 2 (diploid), 3 (polypoid), or NA (unknown). We considered reports of 2*n* = 80 for *Athyrium* and 2*n* = 82 for *Diplazium* as diploid. All other species data were multiples of these numbers (e.g. 2*n* = 160, 2*n* = 123, 2*n* = 164, 2*n* = 205, 2*n* = 246, 2*n* = 328) and were thus scored as polyploids (Table S3). We constructed a variety of models with up to seven cladogenetic rates, denoted as a triplet of values for the ancestral and descendant states: λ_111_, λ_112_, λ_113_, λ_123_, λ_222_, λ_223_, λ_333_, three extinction rates (μ_1_, μ_2_, μ_3_), and four transition rates (*q*_12_, *q*_13_, *q*_21_, *q*_23_) (Fig. 1). We disallowed anagenetic change from polyploidy (3) to diploidy (1, 2) because no diploidization events were reported according to previous cytological studies and our CHROMEVOL analysis (see results). After some preliminary analyses, we settled on six candidate models: 1) a full model with all 13 free parameters; 2) a null model with three parameters, in which diversification is state-independent and anagenetic change is symmetric; 3) a ‘cladogenetic’ model with 11 parameters and a single rate of anagenetic change; 4) an ‘anagenetic’ model with five parameters and a single rate for all speciation and extinction events; 5) an ‘extinction’ model with five parameters and 1 rate each for speciation and anagenetic change; and 6) a ‘cladogenetic-anagenetic’ model with 11 parameters, and a single rate for extinction (Fig. 1; Table S5). Models were run on the whole ingroup clade and separately on each genus, and compared using AIC. Parameters of the best-fit models were estimated using MCMC for 100,000 generations in DIVERSITREE.

**Fig. 1.**
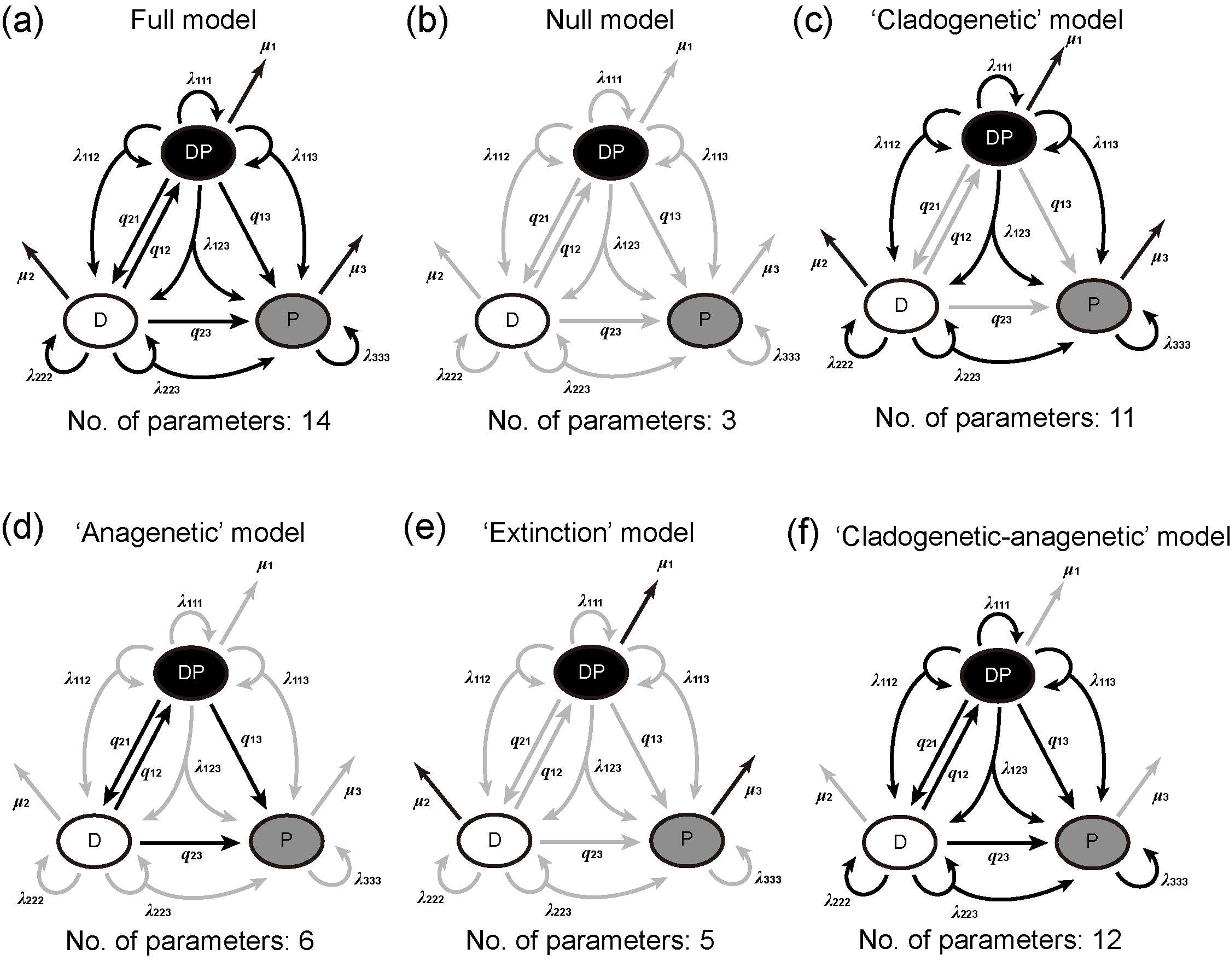
The cladogenetic-state change speciation and extinction (ClaSSE) models with cladogenetic and anagenetic changes. (a) The full model including cladogenetic diversification and anagenetic state change is depicted. In the diagram, lineages presented by both diploid and polyploid populations have state 1, and those with only diploids have state 2, and those with only polyploids have state 3. Each speciation event gives rise to two daughters (as shown by arrows), either in the same state (at rates λ_111_, λ_222_, and λ_333_) or in different states (λ_112_, λ_113_, λ_123_, and λ_223_). Thus, changes in ploidy can occur through the anagenetic (*q*_12_, *q*_13_, *q*_21_, and *q*_23_) or cladogenetic pathway (λ_112_, λ_113_, λ_123_, and λ_223_). Extinction rates in each state are represented by μ_1_, μ_2_ and μ_3_. The rates in gray indicate equal rates, respectively. (b) The null model with equal speciation, extinction and transition rates, respectively. (c) The ‘cladogenetic’ model. (d) The ‘anagenetic’ model. (e) The ‘Extinction’ model. (f) The ‘cladogenetic-anagenetic’ model.

To test the reliability of our ClaSSE analysis given that nearly 40% of the sampled species have unknown ploidy, we simulated 100 trees of the same size as our ingroup clade using the maximum likelihood parameter values of the best-fit model. For each tree we ‘impoverished’ the tip states by randomly setting 40% to the unknown state, NA, and performed model selection using the candidate pool as described above. We also compared the parameters estimated from the impoverished data to their simulation values. All procedures were carried out using DIVERSITREE 0.9-10.

### Reconstruction of polyploidy evolution

To infer the evolutionary history of polyploidy along the phylogeny, we carried out an ancestral state reconstruction using CHROMEVOL 2.0 (Glick & Mayrose, 2014) implemented in RASP 4.0 (Yu *et al.*, 2015). To shorten the calculation time, we used the ‘Auto_run’ option, which automatically optimized all the parameters for chromosome evolution models, including rates of single ascending dysploidy (γ), descending dysploidy (δ), whole-genome duplication (ρ) and demi-polyploidy (μ_d_). After model optimization, 10,000 simulations were performed using both maximum likelihood and Bayesian approaches to summary the final reconstruction result.

## Results

### Divergence time estimation

Our molecular dating analysis yielded age estimates that are broadly congruent with those of previous studies with the variations less than 10 Myr (Schneider *et al.*, 2004; Schuettpelz & Pryer, 2009; Rothfels *et al.*, 2015; Wei *et al.*, 2015; but see Testo & Sundue, 2016 using a different strategy of divergence time estimation based on a time calibrated phylogeny) (Fig. S1). Those that differed tended to be older (e.g. the crown node of the *Diplazium* clade, 47.9 Myr [95% HPD: 41.1–59.4 Myr] in the present study; 41.7 Myr [95% HPD: 33.6–49 Myr] in Wei *et al.*, 2015). This could be due, at least in part, to our much denser sampling of *Athyrium* and *Diplazium* (214 species versus 92 in Wei *et al.*, 2015).

### Diversification analyses: BAMM, QuaSSE and ClaSSE

The BAMM analysis indicated significant increases in net diversification rate in both *Athyrium* and *Diplazium* (Figs 2a and S2; Table S6). In *Athyrium*, two shifts were inferred at 16.86 Myr and 5.02 Myr, with the latter leading to a rate 17 times higher than the root rate (0.044–0.76 events Myr^−1^ per lineage; Fig. 2a). In *Diplazium* one shift was inferred at 23.43 Myr (Fig. 2a). Rates-through-time (RTT) plots revealed contrasting diversification dynamics between *Athyrium* and *Diplazium* following their initial shifts (Fig. 2b). In *Athyrium*, the net diversification rate increased continuously, while in *Diplazium* the rate plateaued to a level about half that of *Athyrium* at the present.

**Fig. 2.**
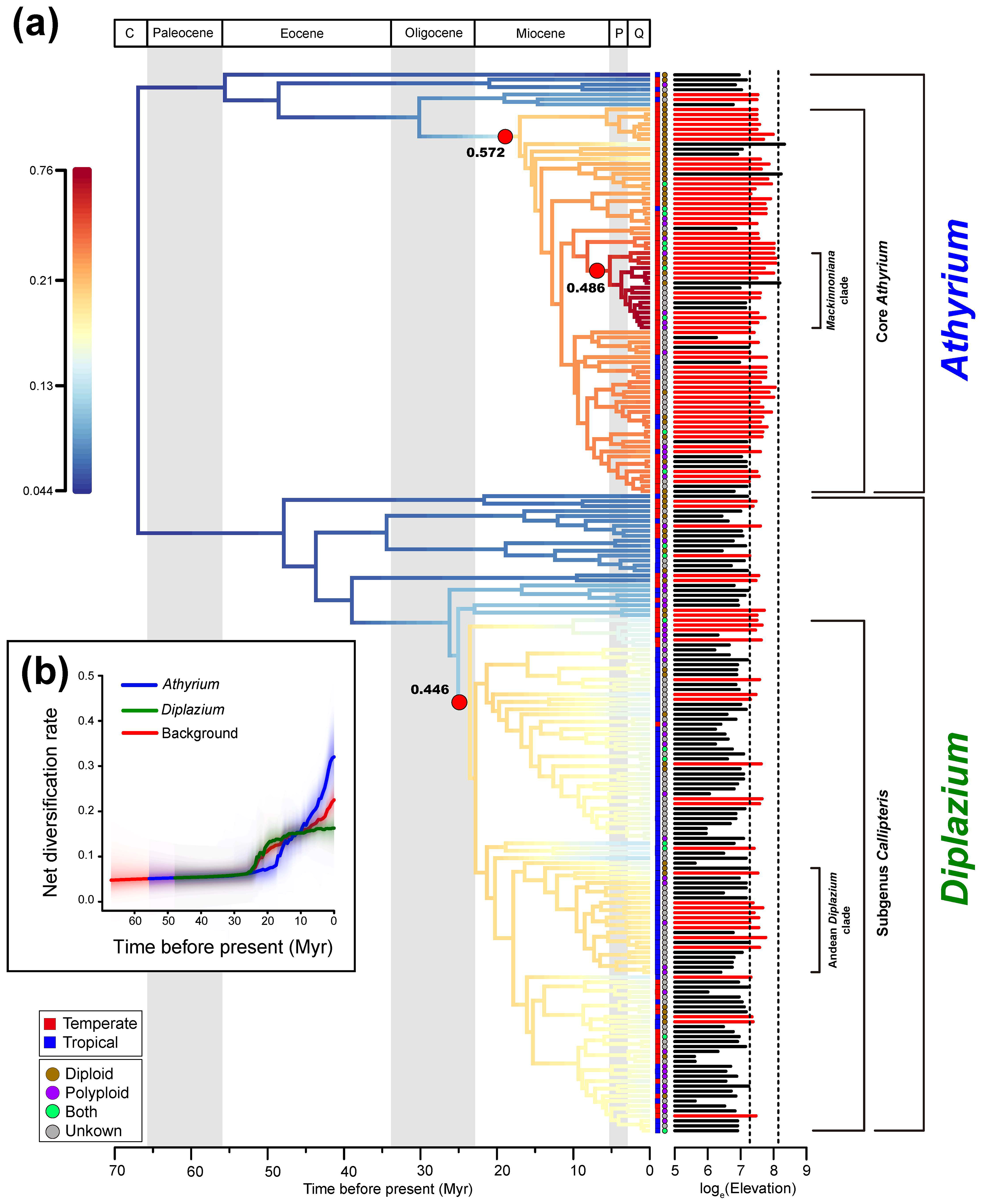
Results of the Bayesian analysis of macro-evolutionary mixtures (BAMM) and quantitative-state speciation and extinction (QuaSSE) analyses. (a) Chronorate plot with branches colored by speciation rate (events Myr^−1^ per lineage) as indicated by the scale bar, representing a summary of the full post-burn-in Markov chain Monte Carlo (MCMC) sample of the BAMM analysis. Red circles indicate the positions of regime shifts in the maximum *a posteriori* (MAP) configuration. Numbers beneath the shifts indicate the marginal probability of a shift occurring along that branch. Log elevation is shown by the horizontal bar for each species. The vertical dashed lines indicate the approximate ranges of altitudes in which elevated speciation rates were inferred by QuaSSE analysis, and extant species whose elevation falls in this range have their data colored red. (b) Rate-through-time plots for speciation rate (events Myr^−1^ per lineage) with 95% confidence interval indicated by yellow shaded areas. Red, the rate across the phylogeny (‘background rate’); blue, the rate of *Athyrium* clade; green, the rate of *Diplazium* clade.

Diversification is associated with elevation itself but not its evolutionary rate. The PGLS regression of the rate of change in elevation on net diversification was nonsignificant (*r*^2^ = 0.286, *P* = 0.379). By contrast, the best-fit QuaSSE model (*wi* = 0.88; Table S4) for the *Athyrium-Diplazium* clade included a unimodal relationship between λ and elevation, with λ highest between 1440 m to 3500 m and constant μ (Fig. 2a). There was a negative directional tendency in *Athyrium* (−0.091) and the entire ingroup clade (−0.086), but a relatively low negative directional tendency in *Diplazium* (−0.0013) (Table 1). Separate analyses of *Athyrium* and *Diplazium* showed differences in their unimodal curves for λ. In *Diplazium* the peak is at a lower rate and elevation, and the curve closely matches the frequency distribution of species by elevation, while in *Athyrium* the peak is higher in rate and elevation, and the curve is shifted right relative to the species’ frequency distribution (Fig. 3). The best-fit model for *Athyrium* includes a negatively valued directional tendency parameter, but the one for *Diplazium* does not (Table 1).

**Table 1.**
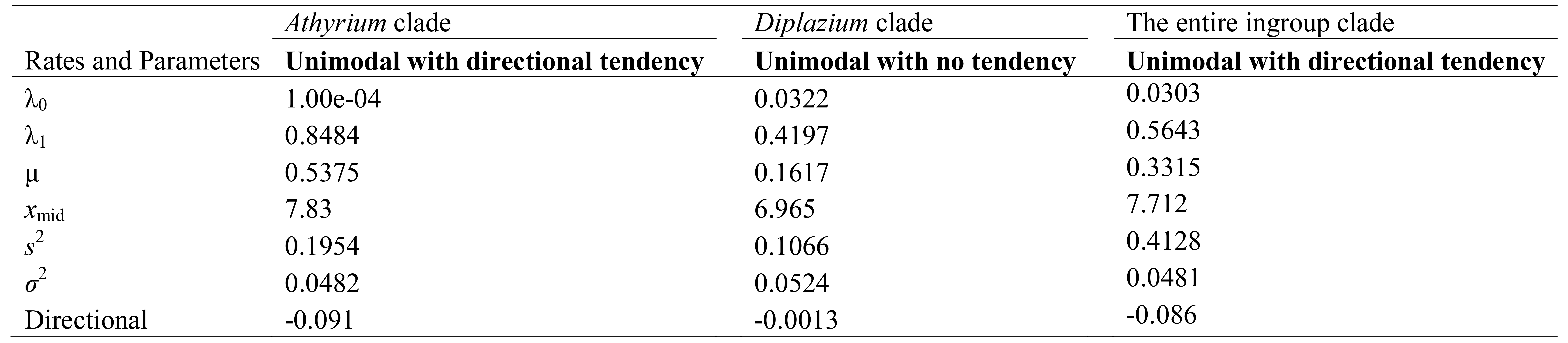
Parameter comparison of the best-fit model of quantitative state speciation and extinction (QuaSSE) analysis based on the entire ingroup clade as well as separately on *Athyrium* and *Diplazium*. λ_0_, rate at lowest values of the log elevation for sigmoid and unimodal models; λ_1_, maximum speciation rate for the unimodal models; *x*_mid_, inflection point of the sigmoid or the place of the maximum for modal models; *s*^2^, width (variance) of the Gaussian function; μ, rate of extinction; σ^2^, Brownian diffusion rate of trait evolution.

**Fig. 3.**
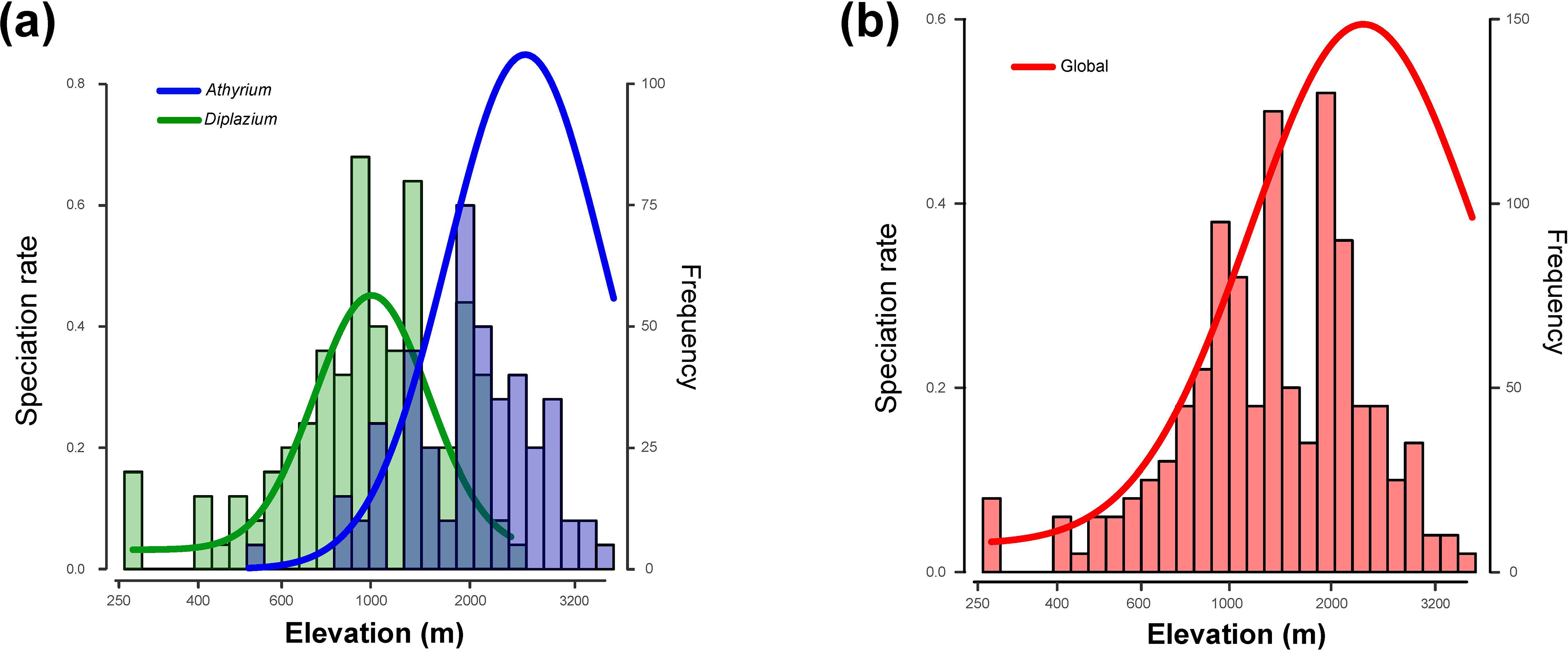
Relationship between the elevation and speciation rates as inferred from QuaSSE analysis, with comparison to the elevational species richness histograms of the *Athyrium* clade, the *Diplazium* clade and the whole phylogeny, respectively. Red, the rate across the phylogeny; blue, the rate of *Athyrium* clade; green, the rate of *Diplazium* clade.

Our ClaSSE analyses yielded strong support for ploidy-dependent diversification dynamics. The ‘cladogenetic’ model, with 10 free parameters for speciation and extinction but no differences in q was selected by AIC (Fig. 4a,b,c,d; Table 2). With this model, rates of speciation in polyploid (λ_3_ = 0.7054 events Myr^−1^ per lineage) and polymorphic species (λ_1_ = 1.388 events Myr^−1^ per lineage) exceed rates for diploids (λ_2_ = 0.1993 events Myr^−1^ per lineage), with the total rate being highest for polymorphic species (Fig. 4a,d; Table 2). Extinction rates are low for diploid and polymorphic species but high for polyploids (Fig. 4c); as a result, the net diversification rate of the polymorphic state is the highest, the rate for diploids is lower but positive, and the rate for polyploids is lowest and slightly negative (Fig. 4e). However, the net accumulation rate of polyploids is highest, followed by diploids and polymorphic species (Fig. 4f). In our simulation analysis of the effect of missing data on model selection, 93% of replicates recovered the correct cladogenetic model (Fig. S3; Table S7), indicating that in this case, model selection is not obviously biased by 40% missing data.

**Fig. 4.**
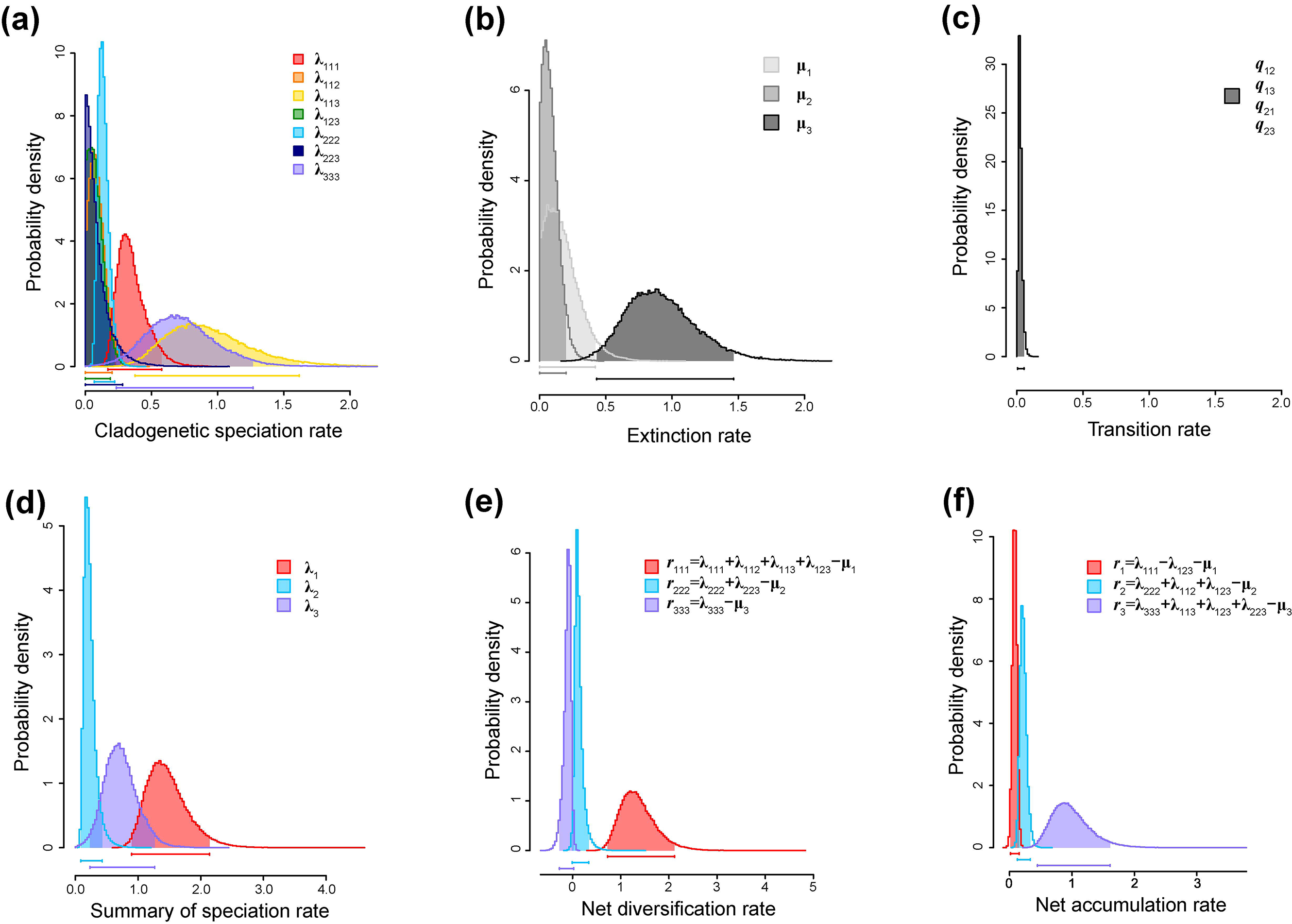
Posterior distributions of macroevolutionary rates under the best-fitting model of the ClaSSE analysis based on the MCC tree, allowing cladogenetic speciation and extinction of diploids and polyploids. (a) The cladogenetic speciation rates of diploid-polyploid mediated speciation events. (b) The extinction rates of three ploidy states. (c) The transition rate of all anagenetic change (equal). (d) Summary of speciation rates of each state. (e) The net diversification rates of ploidy in cladogenetic speciation. (f) The net accumulation rates of ploidy in cladogenetic speciations.

**Table 2.**
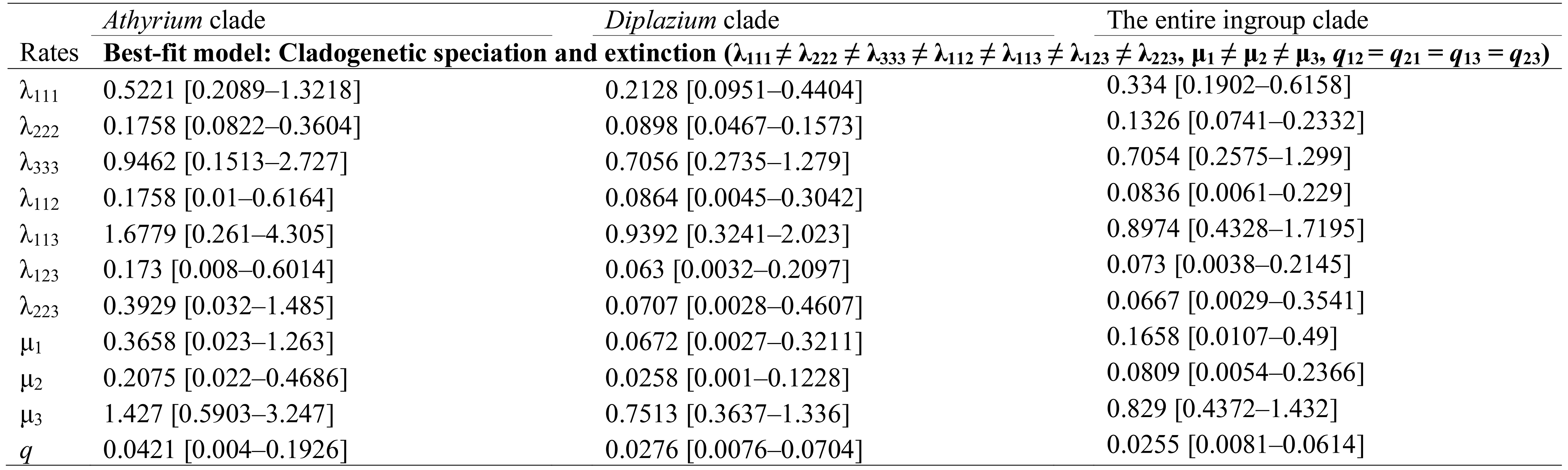
Parameter comparisons of the best-fit model in cladogenetic state change speciation and extinction (ClaSSE) analysis based on the entire ingroup clade as well as separately on *Athyrium* and *Diplazium*. Both containing diploidy and polyploidy is coded as 1; diploidy is coded as 2; polyploidy is coded as 3. λ, speciation rate; μ, rate of extinction; *q*, transition rate between two states. All rates are presented with median values and 95% highest posterior density (HPD) intervals in brackets based on 100 000 MCMC optimization.

Our CHROMEVOL analysis inferred no dysploidization or diploidization events, and at least 15 polyploidization events indicating that all polyploid species in *Athyrium* and *Diplazium* are neopolyploids (Fig. S4).

## Discussion

### Latitudinal contrasts in diversification between temperate *Athyrium* and tropical *Diplazium*

Our dated phylogeny and BAMM analysis showed that species diversity in the primarily temperate genus *Athyrium* has accumulated relatively recently, with an increasing net rate through the Neogene, compared to its primarily tropical sister clade *Diplazium*, which showed dynamics more suggestive of diversity-dependence. Similar temperate-tropical rate contrasts that run counter to the latitudinal diversity gradient have been found in fishes, birds, mammals, and seed plants (Weir & Schluter, 2007; Schluter, 2015; Spriggs *et al.*, 2015), and contradict the hypothesis that higher diversity in the tropics is the result of faster per-lineage diversification (e.g. Mittelbach *et al.*, 2007). The *Athyrium-Diplazium* contrast may reflect how diversification has responded at a coarse scale to waxing and waning ecological opportunities in temperate and tropical habitats, respectively. In this view, the relatively recent and rapid radiation in *Athyrium* was driven by increasing niche availability in expanding temperate habitats, while for *Diplazium*, the contraction of tropical forests had the opposite effect, as global cooling began in the Late Oligocene (Zachos *et al.*, 2001; Morley, 2003). The plateau of net diversification rate in *Diplazium* suggests that the clade is closer to ecological saturation than *Athyrium*, in which the upward trend in rate suggests a clade in the early stages of rapid growth, further from equilibrium (Fig. 2b). These dynamics are consistent with the idea that species diversity is predicted by area, and other proxies for carrying capacity, integrated over time (Fine & Ree, 2006; Jetz & Fine, 2012). In addition, the relatively high turnover rate μ/λ in *Athyrium* (Fig. S5) may be due to more severe climatic oscillations in the temperate zone than in the comparatively stable tropics over the past 20 Myr or so (Tiffney & Manchester, 2001; Morley, 2003), consistent with the latitudinal gradient in turnover found in previous studies (Jablonski *et al.*, 2006; Weir & Schluter, 2007; but see Rabosky *et al.*, 2015). This high rate of species turnover, in turn, suggests a rapid species replacement through non-adaptive evolution in *Athyrium* (Rundell & Price, 2009).

Additional support for the idea that diversification dynamics in *Diplazium* are closer to an equilibrium state than *Athyrium* is provided by our QuaSSE analysis of elevation. In *Diplazium* the inferred response of speciation rate to elevation closely matches the frequency distribution of species along elevation (also is reflected by a low negative directional tendency; Table 1), suggesting a stable state, while in *Athyrium* the rate curve is right-shifted: speciation is fastest at elevations higher than the peak of species richness (Fig. 3). The relative lack of high-elevation species of *Athyrium* could be explained partly by the inferred negative directional tendency in the model – i.e. species originated at high elevations but evolved anagenetically toward lower elevations – but it seems more plausible that *Athyrium* has simply colonized mountains too recently for the number of high-elevation species to reach equilibrium. We would expect such a lag time to be lengthened by the relatively high extinction rate inferred for *Athyrium* versus *Diplazium* (Table 1).

Our analyses of elevation stand in contrast to those in the epiphytic fern family Polypodiaceae, in which faster diversification was positively correlated with the rate of change in elevation, not elevation itself, suggesting a process of adaptive divergence and niche-filling along elevation gradients (Sundue *et al.*, 2015). We were unable to replicate this result from our data, possibly because of insufficiently dense taxon sampling. Nevertheless, our results suggest that diversification is accelerated by occurring in mountains, especially for *Athyrium*. However, it is worth noting that even within *Diplazium*, the highest net diversification was inferred in the lone Andean clade (Fig. 2a). Left open are questions about how and why, such as to what degree is speciation accelerated by non-adaptive processes (such as allopatric genetic drift) versus those involving adaptation? More focused studies of possible mechanisms are needed. One might look for interactions between traits with environmental tolerances and mountain buildings in tropical and temperate zones (Xing & Ree, 2017; Hughes & Atchison, 2015). This has been shown in studies of other plant groups have found that the evolution of traits such as growth form, leaf size, pollination syndromes, and metabolic pathways may be linked to faster diversification in mountains where seasonally cold and arid habitats are dominant (Arakaki *et al.*, 2011; Drummond *et al.*, 2012; Roquet *et al.*, 2013; Schwery *et al.*, 2015; Lagomarsino *et al.*, 2016).

### Polyploidy-driven diversification in the *Athyrium-Diplazium* clade

In ferns, several traits, including growth habit, leaf size, leaf morphology and gametophyte morphology, have been studied for their effect on diversification (Sundue *et al.*, 2015; Ramírez-Barahona *et al.*, 2016; Testo & Sundue, 2018), but polyploidy, despite its ubiquity and precedent in being implicated in diversification (Wood *et al.*, 2009), has thus far escaped intensive quantitative analysis. While the macroevolutionary consequences of polyploidy remain a subject of debate (e.g. Mayrose *et al.*, 2011, 2015; Zhan *et al.*, 2014; Soltis *et al.*, 2014), our inference of rapid turnover with negative net diversification in polyploids (Fig. 3d) is consistent with the ‘extinction-risk’ hypothesis, that neopolyploids suffer deleterious genetic effects (Wright, 1969; Orr, 1990; Ramsey & Schemske, 2002), and thus are often evolutionary dead ends (Mayrose *et al.*, 2011; Arrigo & Baker, 2012). This effect is counterbalanced by the high net diversification rate of polymorphic species (Fig. 4d), which rapidly produce polyploids through cladogenesis events and thereby contribute to polyploids having the highest net accumulation rate (Fig. 4e). Rapid and recurrent neopolyploidy in *Athyrium* and *Diplazium* (Fig. S4) is distinct from *Asplenium*, which is characterized by a dominance of paleopolyploids (Schneider *et al.*, 2017). Taken together, these results imply that although neopolyploids in *Athyrium* and *Diplazium* tend to be short-lived, they play a key role in diversification dynamics (Vamosi *et al.*, 2018).

Infraspecific variation in ploidy is common in plants, and many case studies have examined the origins of polyploidy at the population level, especially in genera of Asteraceae such as *Tragopogon* (Symonds *et al.*, 2010), *Artemisia* (Pellicer *et al.*, 2010), *Helianthus* (Bock *et al.*, 2014), as well as in fern genera *Asplenium* (e.g. Dyer *et al.*, 2012) and *Dryopteris* (e.g. Ekrt & Koutecký, 2016), and *Pteris* (Chao *et al.*, 2012). This study is the first to explicitly model how such polymorphic species may contribute to diversification dynamics at a macroevolutionary scale, and so it seems noteworthy to find that despite occurring at low frequency, their net diversification rates are in fact higher than pure diploids or polyploids. This is largely driven by λ_113_, the rate at which a polymorphic ancestor splits into polymorphic and polyploidy descendants (Fig 4a). Mechanistically, it seems reasonable to predict that this parameter reflects how newly formed polyploids in a population are likely to be reproductively isolated from their diploid progenitors, and thus form a new lineage. Infraspecific polyploidy can predispose species to be better adapted to harsher conditions in novel environments (Levin, 1983; Ramsey & Schemske, 2002; te Beest *et al.*, 2012). These results could be framed as a ‘ploidy-polymorphism compensation’ hypothesis, wherein polymorphic species play a central role in the maintenance of neopolyploid diversity. The *Athyrium-Diplazium* clade is not likely to be unique in supporting this hypothesis; we predict that other lineages such as *Asplenium* (Aspleniaceae) and *Dryopteris* (Dryopteridaceae) and *Pteris* (Pteridaceae), in which autopolyploidy, hybridization and apomixis are prevalent (Chao *et al.*, 2012; H.M. Liu *et al.*, 2012), may have similar compensatory dynamics mitigating the high extinction risk of polyploids.

Why might polyploids have high speciation rates (Fig. 4a)? It is possible that some factors (a novel physical trait or a property of the environment) associated with polyploidization may play a role (Zhan *et al.*, 2016; Vamosi *et al.*, 2018). A few case studies have linked polyploidization to trait evolution and diversification (Schranz *et al.*, 2011; Tank *et al.*, 2015). In ferns, polyploids may facilitate long-distance dispersal via tolerance of inbreeding, i.e. gametophytic selfing, increasing the success rate of single-spore colonization (e.g. Tryon, 1985; Testo *et al.*, 2015; Sessa *et al.*, 2016). This in turn might favor colonization of different ecological niches, and thus increase the probability for peripatric speciation. In *Deparia*, the species-rich genus in Athyriaceae that is sister to *Athyrium-Diplazaium*, polyploid species are characterized by increased dispersal abilities and greater range expansion than sexual diploids (Kuo *et al.*, 2016). This seems to be true as well for *Athyrium* and *Diplazium*, in which polyploid species are often found more broadly distributed than their diploid relatives (Tryon & Tryon, 1982; Takamiya *et al.*, 1999, 2000; Takamiya & Ohta, 2001; Bir & Verma, 2010). It thus seems necessary for future studies to target other traits potentially linked polyploidy that could enhance dispersal ability.

## Acknowledgements

We thank Qin Li, Matthew Nelsen, Shrabya Timsina, and Charles Bell for helpful discussions and Jen-Pan Huang for technical assistance and advice about ClaSSE. We thank Harald Schneider, Hong-Mei Liu, Yue-Hong Yan, Hui Shang, Atsushi Ebihara, Michael Kessler, Marcus Lehnert, Peter Hovenkamp, Ronald Viane, Sabine Hennequin and Claudine Mynssen for providing valuable materials. This study was supported by grants from the National Science Foundation of China Grant (No. 31600175), the Beijing Natural Science Foundation Grant (No. 5162020) and the Visiting Scholar of Bureau of Personnel Chinese Academy of Sciences to the Field Museum awarded to R.W.

## Author contributions

X.-C.Z. and R.W. conceived the idea of this study; R.W. performed the experiments; M.A.S. provided important Neotropical materials; R.W. and R.H.R. analyzed data; and R.W., R.H.R., and M.A.S. wrote the paper, with significant contribution from X.-C.Z.

## Supporting Information

Additional Supporting Information may be found online in the Supporting Information tab for this article:

**Fig. S1** Chronogram of the *Athyrium-Diplazium* clade with four calibration points.

**Fig. S2** Positions of regime shifts in the maximum a posteriori configuration inferred from the Bayesian analysis of macro-evolutionary mixtures (BAMM).

**Fig. S3** Rate simulations and their estimates of confidence intervals for each of the states under 100 replicates of simulated trees and random assignment of ‘NA’ to tip states in ClaSSE.

**Fig. S4** CHROMEVOL inferences for the *Athyrium-Diplazium* clade.

**Fig. S5** Rate-through-time plot of rates of speciation, extinction and species turnover (termed ‘μ/λ’) obtained from BAMM.

**Table S1** Species sampling fraction used in macroevolution analyses.

**Table S2** Taxa, voucher information and GenBank accession numbers of specimens used in this study.

**Table S3** Traits information and coding in QuaSSE, CHROMEVOL and ClaSSE.

**Table S4** Model selection and results of QuaSSE on the entire ingroup clade as well as separately on *Athyrium* and *Diplazium*.

**Table S5** Model selection and results of ClaSSE based on the entire ingroup clade as well as separately on *Athyrium* and *Diplazium*.

**Table S6** Bayes factor of different diversification rate models fitted on the *Athyrium-Diplazium* dataset.

**Table S7** ClaSSE model simulation based on 100 simulated trees with random assignment of missing information to tips.

